# Biological, Molecular and Phiysiological Characterization of Four *Soybean mosaic virus* Isolates Present in Argentine Soybean Crops

**DOI:** 10.1101/2021.06.10.447356

**Authors:** M Maugeri Suarez, M Rodríguez, N Bejerman, I. G Laguna, P Rodríguez Pardina

## Abstract

*Soybean mosaic virus* (SMV) causes systemic infections in soybean plants, leading to chlorotic mosaic and producing significant yield losses. The virus is widely distributed in all soybean production areas in the world. In Argentina, three geographical isolates were identified: Marcos Juárez (MJ), Manfredi (M), and North Western Argentina (NOA), and another isolate named “Planta Vinosa” (PV), which causes severe necrosis symptoms in some cultivars. Here, the biological, molecular and physiological characterization of these isolates was performed for the first time. Three of the four isolates showed a low genetic divergence in the evaluated genes (P1, CI and CP). Although SMV-NOA and SMV-PV had high homology at the sequence level, they showed wide differences in pathogenicity, seed mottling and the ability of transmission by seeds or aphids, as well as in physiological effects. SMV-NOA caused early alterations (before symptom appearance, BS) in ΦPSII and MDA content in leaves with respect to the other isolates. After the appearance of macroscopic symptoms (late symptoms, LS), SMV-M caused a significant increase in the content of MDA, total soluble sugars, and starch with respect to the other isolates. Thus, early alterations of ΦPSII and soluble sugars might have an impact on late viral symptoms. Likewise, SMV-MJ developed more severe symptoms in the susceptible Davis cultivar than in DM 4800. Therefore, our results show differences in genome, biological properties and physiological effects among SMV isolates as well as different interactions of SMV-MJ with two soybean cultivars.

## INTRODUCTION

Soybean [*Glycine max* (L) Merr.] is one of the most important legume crops and a source of edible oil and proteins. Argentina is the third world soybean producer, with 84% of the production being exported as grain, flour, oil or biodiesel (FAOSTAT 2019). Extensive and intensive production of soybean with little genetic diversity is particularly vulnerable to attack by pathogens that can reduce yield and seed quality, and even devastate big cultivation areas. Virus diseases of soybean have become increasingly prevalent, affecting this crop worldwide. *Soybean mosaic virus* (SMV) is recognized as the most serious, long-standing problem in many soybean-producing areas in the world (Wang, 2009; Cui et al., 2011). Infection by SMV usually causes yield losses ranging between 35 and 50%, with estimates of 50–100% in severe outbreaks (Arif & Hassan, 2002; Liao et al., 2002).

SMV produces variable symptoms, from small and sometimes almost unnoticeable chlorotic spots, to large chlorotic areas. Other possible symptoms include mosaic, vein clearing, blistering, leaflet deformation and internode shortening. When plants are infected with severe virus strains, the virus can induce necrotic areas in petioles, stems and leaves (Hajimorad et al., 2018). SMV also induces several types of seed mottling, with the most common one being “hilum bleeding”, caused by the spread of the hilum color towards the seed coat. SMV-infected seeds can result in infected plants that serve as the initial inoculum with later infections resulting from aphid transmission. Seed infection can be as high as 75%, depending on the soybean cultivar and the virus strain, but is usually less than 5% (Rupe & Luttrell, 2008; Sweets, L, 2011).

SMV is a member of the genus *Potyvirus*, in the *Potyviridae* family. It has a monopartite single strand, positive-sense RNA genome that encodes a large polyprotein of about 350 kDa. This polyprotein is cleaved to yield at least 11 proteins: potyvirus 1 (P1), helper component proteinase (HC-Pro), potyvirus 3 (P3), PIPO, 6 kinase 1 (6K1), cylindrical inclusion (CI), 6 kinase 2 (6K2), nuclear inclusion a-viral protein genome linked (Nia-Vpg), nuclear protein a-protease (NIa-Pro), nuclear inclusion b (NIb) and coat protein (CP) (R.-H. Wen & Hajimorad, 2010). Several SMV isolates were classified into different strains based on their differential response in susceptible and resistant soybean cultivars (Buzzell & Tu, 1984; Cho & Goodman, 1982; S. M. Lim, 1985; Pu et al., 1982; Zhan et al., 2006). Different types of responses of susceptible and resistant cultivars are the result of specific interactions between the soybean *R* gene product and the virus avirulence (*Avr*) gene product. At least three independent loci (*Rsv1, Rsv2 and Rsv4*) in the United States and several *Rsc* loci in China conferring resistance to different SMV strains have been reported (Liu et al., 2016). Identification of SMV strains is very important for both soybean cultivation and breeding. The method based on the pathogenicity has been widely used; however, this method is laborious and time-consuming. Therefore, genomic sequences have also been used to differentiate SMV strains in recent years. In this sense, one of the most variable and informative proteins to compare strains is P1 (Domier et al., 2003; W. S. Lim et al., 2003).

Studying the molecular variability and genetic structure of viruses helps to provide understanding of their molecular evolutionary history in relation to virulence, dispersion and emergence of new epidemics (Seo et al., 2009). These studies focused mainly on phylogenetic relationships between virus isolates, because most of the viruses are constantly evolving through genetic exchanges (recombination), as well as accumulation of mutations (Choi et al., 2005; Gagarinova et al., 2008; Saruta et al., 2005). Due to the rapid evolution in avirulence/effector genes, the resistance conditioned by genes will be quickly overcome and it is important to generate strategies for the management of viral diseases that are sustainable over time (Liu et al., 2016). In this context, there is a significant demand to identify plant factors involved in defense responses to pathogens that can facilitate the design of new sustainable tolerance/resistance strategies against SMV. Therefore, it is necessary to know the impact of viruses on plant physiology, as well as the mechanisms and processes involved in the infection.

A compatible plant-virus interaction causes deleterious systemic effects on plants because viruses have the capacity to reprogram the plant metabolism to their own benefit (Andreola et al., 2019, Zanini et al., 2021). Reprogramming includes suppression of plant defense responses, reallocation of photoassimilates, redox imbalance, reduced photosynthesis and induced senescence (Loebenstain & Carr, 2006, Andreola et al., 2019).

In Argentina, the isolates G1, G5, G6 (from USA), and MS1 and MS2 (Brazil) were detected (Truol & Laguna, 1992). In addition, three geographic isolates of this virus, Marcos Juárez (MJ), Manfredi (M), and northwestern Argentina (NOA), and an isolate called “PlantaVinosa” (PV), which that causes severe necrotic symptoms in some cultivars, were collected for further characterization. The aim of the present study was to perform the biological, physiological and molecular characterization of the latter four isolates. Therefore, combined information about genetic and physiological alterations in the SMV-soybean interaction is provided for the first time.

## MATERIAL AND METHOD

### Inoculum source

Plants with SMV symptoms were collected from four soybean-production areas of three provinces of Argentina: Marcos Juárez (MJ) and Manfredi (M) from Córdoba province, Salta (NOA) and Santa Fe. The isolate detected in Santa Fe causes severe necrotic symptoms in some cultivars and was named “Planta Vinosa” (PV), due to the reddish color observed on stems and petioles, similar to the color of red wine. The inoculums were multiplied in soybean cv DM 4800 and Davis, through mechanical transmissions, with 0.05 M, pH 7.6 potassium phosphate buffer. Soybean plants were grown under controlled conditions: 25 ± 2 °C and a 16:8 light: dark photoperiod (250 μmoL photon. m^-2^. sec^-1^) and 65% humidity.

### Biological characterization

#### Pathogenicity test

Once multiplied, the four isolates were mechanically transmitted to a group of differential cultivars (Table 1), as suggested by several authors (Almeida, 1981; Cho & Goodman, 1982; Shigemori, 1991). The 10 inoculated plants per cultivar/isolate were maintained under greenhouse conditions (25 °C ± 2) until the onset of symptoms. Infection was confirmed using PTA ELISA (Converse & Martin, 1990), through the analysis of the last developed trifoliate leaf. Systemic and local symptoms were recorded.

**Table 1.**
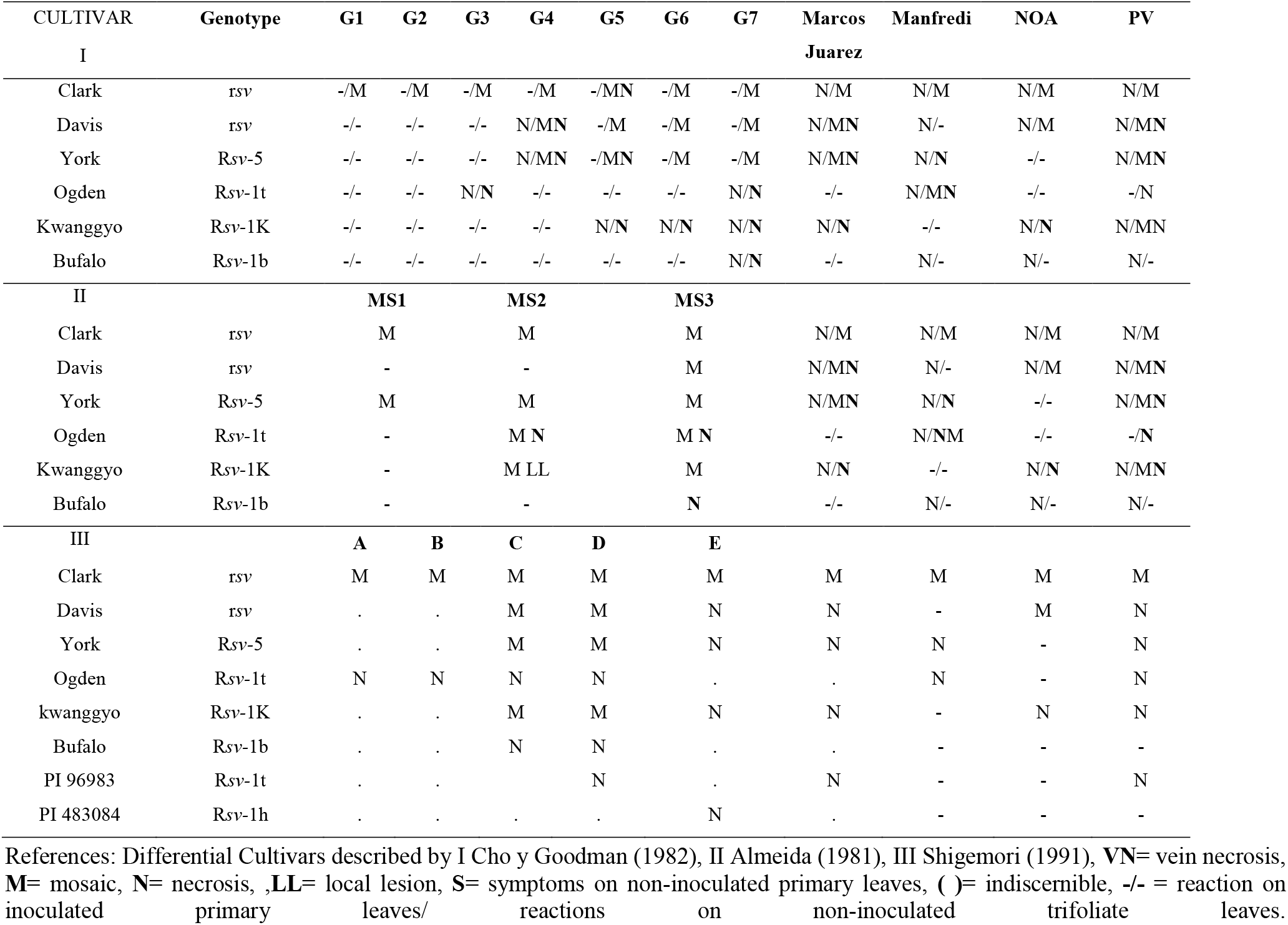
Reactions of differential soybean varieties to four soybean mosaic virus isolates from Argentina

#### Aphid transmission

Colonies of *Myzus persicae* Sulzer were bred on *Ipomea setosa* Nil. Two trials were performed, one using two aphids per plant and the other only one aphid per plant. For transmission, aphids were starved for 3 to 4 hours, then allowed to feed on soybean plants infected with the different SMV isolates for a maximum period of one minute (acquisition); then they were allowed to feed on healthy soybean plants of Forrest cultivar (10 plants/isolate) for approximately 18 hours. The plants were maintained under greenhouse conditions (25 °C ± 2) until the evaluation of transmission through visual symptoms and PTA-ELISA, following a previously described protocol (Converse & Martin, 1990)

#### Seed transmission

Twenty soybean plants of Forrest cultivar per studied isolate (MJ, M NOA, and PV) were mechanically inoculated The inoculated plants were maintained under greenhouse conditions (25 ± 2 °C) until maturity, when pots were harvested, and the percentage and degree of mottled seeds was estimated. To evaluate seed transmission, all the harvested seeds were sown in individual terrines, and seedlings were analyzed by PTA ELISA using the first trifoliate leaf.

#### Molecular characterization

Total RNA was extracted from approximately 200 mg of infected leaves using the Trizol reagent method (Chomczynski & Sacchi, 1987). The obtained RNA was quantified using the nanodrop® ND-1000 Spectrophotometer.

Fragments corresponding to the CI and P1 genomic regions were amplified by RT-PCR, using the sets of primers described by Kim et al. (2004) and Sherepitko et al. (2011) (Table 2). RT-PCR was performed with the Access RT-PCR System (Promega Corporation Madison WI USA), using as template 1 μg of RNA of the different isolates. RT-PCR conditions for the CI segment were as follows: cDNA synthesis at 48 °C for 45 min., 2 min at 94 °C, followed by 40 cycles of 30 sec at 94 °C, 1 min at 60 °C, and 2 min at 68 °C, with a final extension of 7 min at 68 °C. To amplify the P1 fragments, thermocycling was programmed as follows: 48 °C for 45 min., 2 min at 94 °C and 35 cycles of 30 s at 94 °C, 30 sec at 55 °C, 1 min. at 68 °C, and the last extension of 10 min. at 68 °C.

**Table 2.**
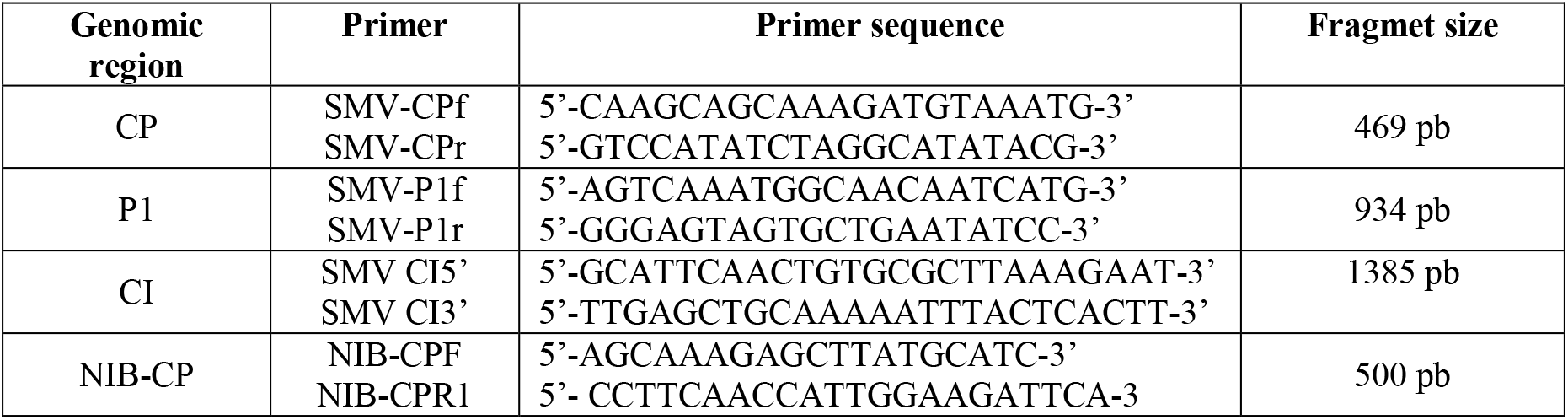
Primers used to amplify the CP, P1 and CI genomic regions

Two pairs of primers, CP and NIB-CP (Table 2), were used for the amplification of the complete CP coding region. The RT-PCR for the CP segment was carried out with the same reaction mix and conditions as those used for P1. For the NIB-CP fragment HotStartTaq Master Mix Kit (Qiagen) was used, and the RT-PCR conditions were as follows: 15 min at 95 °C and 40 cycles of 30 sec at 95 °C, 1 min at 53 °C, 1 min at 72 °C, and the last extension of 10 min at 72 °C. The amplified products were purified with the DNA cleaning and concentrator kit (Zymo Research CA, USA), and sequenced at the Genomic Unit of the Biotechnology Institute-INTA (Argentina). Once obtained, the sequences of P1, CI and CP were assembled with the Seqman tool (DNASTAR Inc. Madison, WI, USA).

The sequences of each isolate were subsequently compared with each other and with those of other SMV isolates, available at the National Center of Biotechnology Information (NCBI - http://www.ncbi.nlm.nih.gov), using the Blastn algorithm (http://www.ncbi.nlm.nih.gov/BLAST) (Altschul et al., 1990). Sequence homology was analyzed with the LASERGENE (DNASTAR Inc. Madison, WI, USA) program. Multiple alignments were performed with *Clustal W* (http://www.justbio.com). Maximum Likelihood (ML) phylogenetic trees were constructed with MEGA X 5.2 program employing the X as best-fit model with 1000 bootstrap iterations (Kamur et al., 2018).RDP, GENECONV, MaxChi, BOOTSCAN, Chimaera, 3Seq and SISCAN methods implemented in the RDP4 (Recombination Detection Program v.4.82) program (Martin et al. 2015) were used to detect recombination events between the different isolates under study. Only those events detected by at least three methods were considered positive.

### Physiological parameters

#### Infection with SMV

*Glycine max* cv. DM 4800 or Davies plants were infected with M, MJ, NOA and PV isolates. Symptomatic leaves were used to prepare the infected extract with 0.05 M, pH7.6 potassium phosphate buffer. SMV infection was performed at 7 days post-germination (dpg) by mechanical damage with carborundum Mesh 600 in the first pair of unifoliate true leaves. The plant was in VC (cotyledon) stage. To evaluate systemically infected leaves, samples were always taken from the first trifoliate leaf at 4 days post-inoculation (dpi) (before symptom expression, BS), and 12 dpi (late symptom expression, LS).

#### Mock infection

mechanical damage with carborundum Mesh 600 was induced with 0.05 M, pH7.6 potassium phosphate buffer.

### Growth parameters

A total of 12 plants per treatment were harvested at the end of the experiment (12 days dpi). To determine Fresh weight (FW) and Dry weight (DW), aboveground tissues were individually harvested. For DW measurements, samples were oven-dried at 80 °C until constant weight was reached. Leaf area was calculated from scanned images of plants 4 and 12 dpi, using Image Pro Plus ver. 4.5.0.29 for Windows 98/NT/2000 image analysis software.

### Chlorophyll fluorescence

Quantum efficiency of PSII photochemistry under ambient light conditions (250 μmoL photon m^-2^ sec^-1^, 25 ± 2 °C) (ΦPSII) was measured using a pulse amplitude modulated fluorometer (FMS2, Hansatech Instruments, Pentney King’s Lynn, UK). Furthermore, leaves were dark-adapted using leaf clips for at least 30 min in order to allow full oxidation of the reaction centers (RC). Then, an actinic 1-sec light pulse of 3500 μmol photons m^−1^ sec^−1^ was applied to reach the maximum fluorescence emission in order to measure Fv/Fm.

### Lipid peroxidation

Lipid peroxidation levels (determined as thiobarbituric acid reactive substances (TBARS)) were measured in the first trifoliate leaf, according to Heath & Packer (1968). The samples were homogenized using a mortar and pestle under liquid nitrogen and thawed in 3% (v/v) trichloroacetic acid (TCA) and centrifuged at 13,000 x *g*, 4 °C during 15 min. A fraction (100 μL) of the sample was mixed with 100 μL of 20% TCA + 0.5% thiobarbituric acid (TBA) and incubated at 90 °C for 20 min; then the samples were rapidly cooled on ice. The mixture was centrifuged at 13,000 x *g* for 10 min. The supernatant was immediately measured by spectrophotometer read at 532 nm and 600 nm absorbance.

### Total soluble sugars and starch

Extracts were obtained following Guan & Janes (1991); 2 g of frozen tissue were ground in 2 ml buffer containing 50 mM HEPES-KOH (pH 8.3), 2mM EDTA, 2mM EGTA, 1mM MgCl^2^, 1mM MnCl^2^, and 2mM dithiothreitol (DTT). The extract was centrifuged at 15,000 rpm at 4 °C for 15 min and the supernatant was used for soluble sugar determination. Soluble sugars were measured with anthrone reagent (Fales, 1951) using sucrose as standard. Starch was determined in the pellet from reducing sugars released after hydrolysis with α amyloglucosidase, (Schneb & Somers, 1944) using glucose as a standard.

### Serological virus detection

Soluble proteins were extracted in coating buffer (Na_2_CO_3_/ NaHCO_3_), pH 9.6, and quantified according to Bradford (1976) without SDS. SMV infection was detected by enzyme-linked immunosorbent assays (PTA-ELISA) using 5 μg of protein per well (Clark and Adams 1977) with anti-SMV-IgG. Polyclonal SMV antiserum. Bovine serum albumin was used as standard for calibration curves. In all cases, six healthy samples and one SMV positive sample per plate were used as controls. Reactions were quantified in Thermo Labsystem MultisKan MS spectrophotometer and samples were considered positive when Abs405 was greater than 0.100 or the mean of healthy controls plus three times the standard deviation (cut-off). Finally, the relative virus concentration was calculated through the A405 of each sample/cut off.

### Statistical analysis

The data obtained were subjected to a parametric analysis of variance (ANOVA), for which the assumptions of Normality and Homogeneity of variances for each variable used were tested. Significant differences (p <0.05) between treatments were evaluated using a DGC multiple range test. All these analyses were carried out with the InfoStat 2015 program (http://www.infostat.com.ar). Statistical analysis were made between treatments, and the values were expressed relative to control.

## RESULTS

### 1. Biological characterization: Pathogenicity tests, aphids and seed transmission

The pathogenicity test performed to characterize the four isolates under study did not allow us to group them with any of the strains previously described by other research groups. The phenotypic severity of the isolates showed differences, with SMV-PV isolate being the most severe one, since it produced mosaic symptoms only in the susceptible cultivar Clark, and caused symptoms of systemic necrosis in the other cultivars, except in Buffalo and PI 483084. On the other hand, the mildest isolate turned out to be NOA, which produced mosaic symptoms in Clark and Davis cultivars, and systemic necrotic symptoms only in the Kwanggyo cultivar (Table 1).

The percentages of transmission by aphids for each isolate, detected by PTA-ELISA, were proportionally similar in both trials (Table 3). The SMV-M, -MJ and -NOA isolates had similarly high percentage of aphid transmission (61%-72%), whereas SMV-PV presented very low transmission capacity (12.5%). Seven days after transmission, all the inoculated plants presented symptoms, such as necrotic and chlorotic local lesions, chlorotic spots and mosaic.

**Table 3.** Transmissibility (%) of soybean mosaic virus Argentine isolates by *Myzus persicae*

|  | M | NOA | MJ | PV |
| --- | --- | --- | --- | --- |
| **2 aphid per plant** | **72** | **67** | **61** | **12,5** |
| **1 aphid per plant** | **37,5** | **38** | **20** | **10** |

Seeds from the Forrest cultivar originated from plants infected with SMV-M isolate presented the highest percentage of mottling (62%) and the highest rate of seed transmission (13%) (Table 4).

**Table 4.** Seed transmission and mottling by different soybean mosaic virus isolates in Forrest cultivar.

| Isolate | % of not spotted seeds | % spotted seeds | Mottling severity* | % seed transmission (DAS-ELISA) |
| --- | --- | --- | --- | --- |
| **Planta Vinosa** | 65 | 35 | 1-2 | 7 |
| **Marcos Juarez** | 58 | 35 | 1-2 | 7 |
| **NOA** | 60 | 39 | 1-3 | 10 |
| **Manfredi** | 38 | 62 | 1-4 | 13 |

\* Mottling severity 1: Mottling covers less than 20% of the seed surface 2: Mottling covers between 20 and 40% % of the seed surface 3: Mottling covers between 40 and 60% % of the seed surface 4: Mottling covers more than 60% of the seed surface

### 2. Phylogenetic characterization and recombination analysis of the SMV isolates

The complete nucleotide sequences of all the evaluated segments/isolates were deposited in the GenBank database. Accession numbers are listed in Table 5. The percentages of similarity and divergence among isolates for each segment are presented in Table 6. SMV-NOA and -PV isolates showed a great similarity (97.5-99.6%) in all the analyzed sequences, whereas a notable divergence (30.6-32.6%) was observed between the above mentioned isolates and SMV-M and -MJ isolates in the P1 segment.

**Table 5.** GenBank Accession numbers of three regions of Soybean mosaic virus genome, corresponding to four different Argentine isolates.

| Soybean mosaic virus region | Isolate | GenBank Accession number |
| --- | --- | --- |
| P1 | M | MH746624 |
| P1 | MJ | MH763836 |
| P1 | NOA | MH795799 |
| P1 | PV | MH785076 |
| CI | M | MH672689 |
| CI | MJ | MH683726. |
| CI | NOA | MH678613. |
| CI | PV | MH688059 |
| Nib-CP | M | MW187865 |
| Nib-CP | MJ | MW187866 |
| Nib-CP | NOA | MW187867 |
| Nib-CP | PV | MW187868 |

**Table 6.** Percentages of similarity and divergence between the four soybean mosaic virus isolates

| Segment | Isolates | Similarity (%) |  |  | Divergence (%) |  |  |
| --- | --- | --- | --- | --- | --- | --- | --- |
| **P1** |  | MJ | NOA | PV | MJ | NOA | PV |
|  | M | 95.9 | 53.1 | 53.6 | 4.2 | **31.4** | **30.6** |
|  | MJ | - | 51.1 | 51.4 | - | **32.3** | **31.9** |
|  | NOA | - | - | **99.1** | - | - | 0.9 |
| **CI** | M | 90.2 | 97.8 | 97.7 | 10.8 | 2.2 | 2.3 |
|  | MJ | - | 91 | 91 | - | 9.8 | 9.8 |
|  | NOA | - | - | **99.5** | - | - | 0.6 |
| **CP** | M | 92.8 | 94 | 95.2 | 7.8 | 6.4 | 5 |
|  | MJ | - | 96.7 | 94.2 | - | 3.4 | 6 |
|  | NOA | - | - | **97.5** | - | - | 2.5 |

The phylogenetic trees are shown in Fig. 1. SMV-NOA and -PV isolates were closely related in CP, P1 and CI sequences, and were associated with the P1 segment of the LJZ010 isolate (China). SMV-M isolate grouped with the TNP strain (USA) in the analysis of fragments P1 and CI, and with strains TNP, G3 and G1 in the analysis of the CP segment. The SMV-MJ isolate was related to the isolates/races WS101, G6, WS32 and G5, and to G5H (South Korea) for the P1 and CP fragments.

**Fig. 1.**
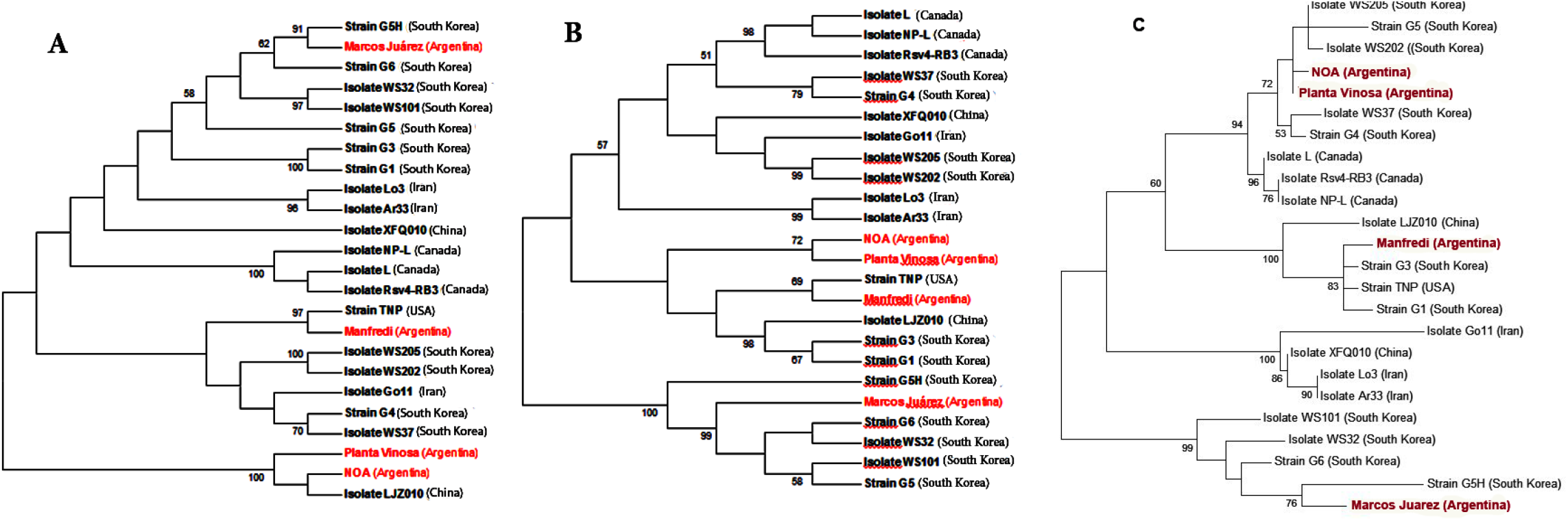
Phylogenetic trees based on the nucleotide sequences of P1 (a). CI (b) and CP (c) segments of MJ. PV. NOA and M. Soybean mosaic virus isolates and other selected soybean mosaic virus strains. Bootstrap values (1000 replicates) are indicated at nodes GeneBank accessions of the soybean mosaic virus strains are: **TNP**: HQ845735; **Rsv4-RB3**: JN416770; **L**: EU871724; **NP-L**: HQ166266. **WS32**: FJ640954; **WS37**: FJ640955; **WS101**: FJ640957, **WS202**: FJ640974; **WS205**; FJ640975; **G1**: FJ640977. **G3**: FJ640978; **G4**:FJ640979; **G5**:AY294044; **G6**:AF242845; **G5H**:FJ807701; **Go11**:KF135491; **Lo3:**KF135490**; Ar33**:KF297335; **LJZ010**:KP710866; **XFQ010**:KP710874.

According to the recombination analyses, SMV-NOA, -PV and -M isolates may have arisen by recombination (Figs 2-3). The P1 segments of SMV-NOA and -PV isolates presented the same recombination event, with exchange points being nucleotides 690 and 1128 approximately for SMV-NOA, and 690 and 1185 for SMV-PV. LJZ010 and G4 were detected as the major and minor parental sequences, respectively. On the other hand, in the analysis of the CI segment, SMV-M isolate, as the NPT strain, would be recombinant between G4 (parent major) and G3 (parent minor), and the exchange points for SMV-M were nucleotides 1560 and 1902. The analysis of the CP fragment showed no recombination events.

**Fig. 2.**
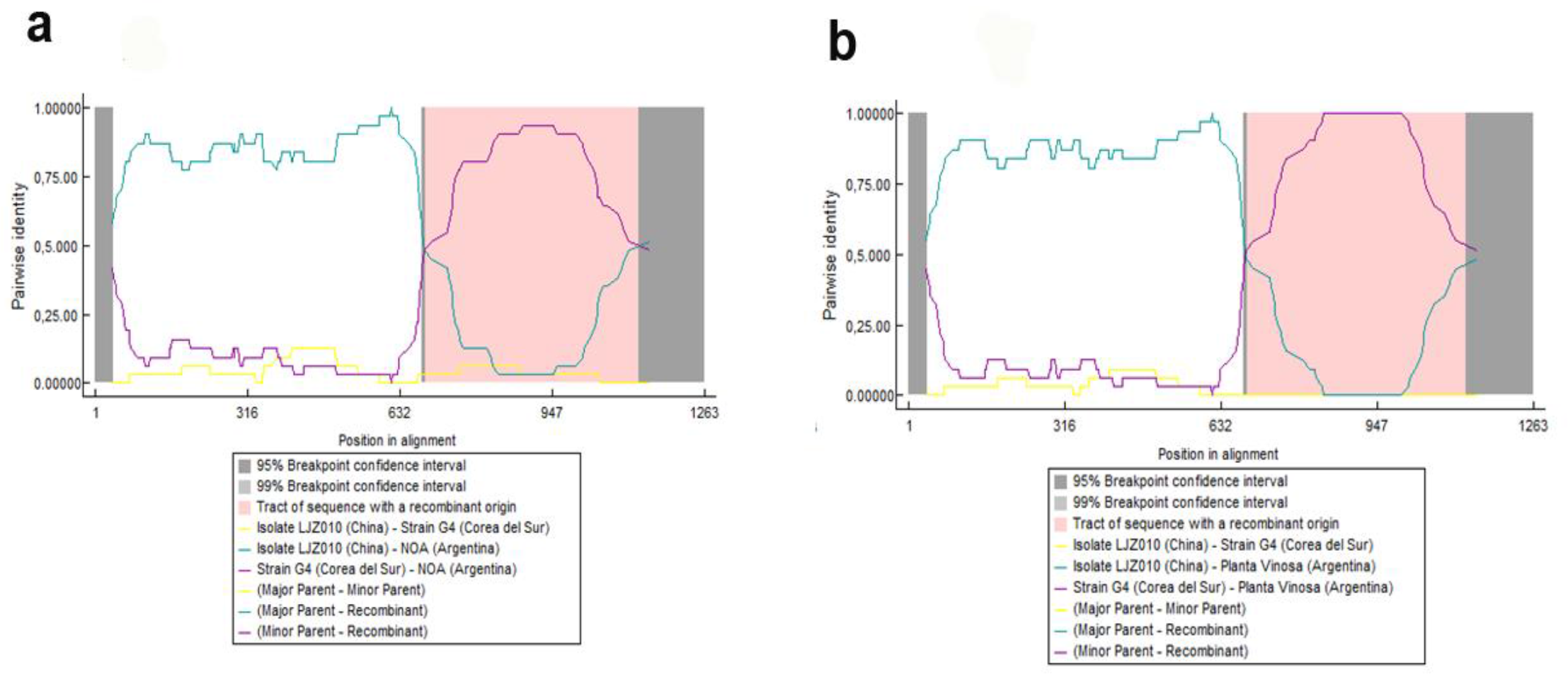
DNA Recombination events detected in the P1 segment of NOA (a) and PV (b) isolate with parental LJZ010 (light blue) and G4 (violet); the corresponding breakpoints are included.

**Fig. 3.**
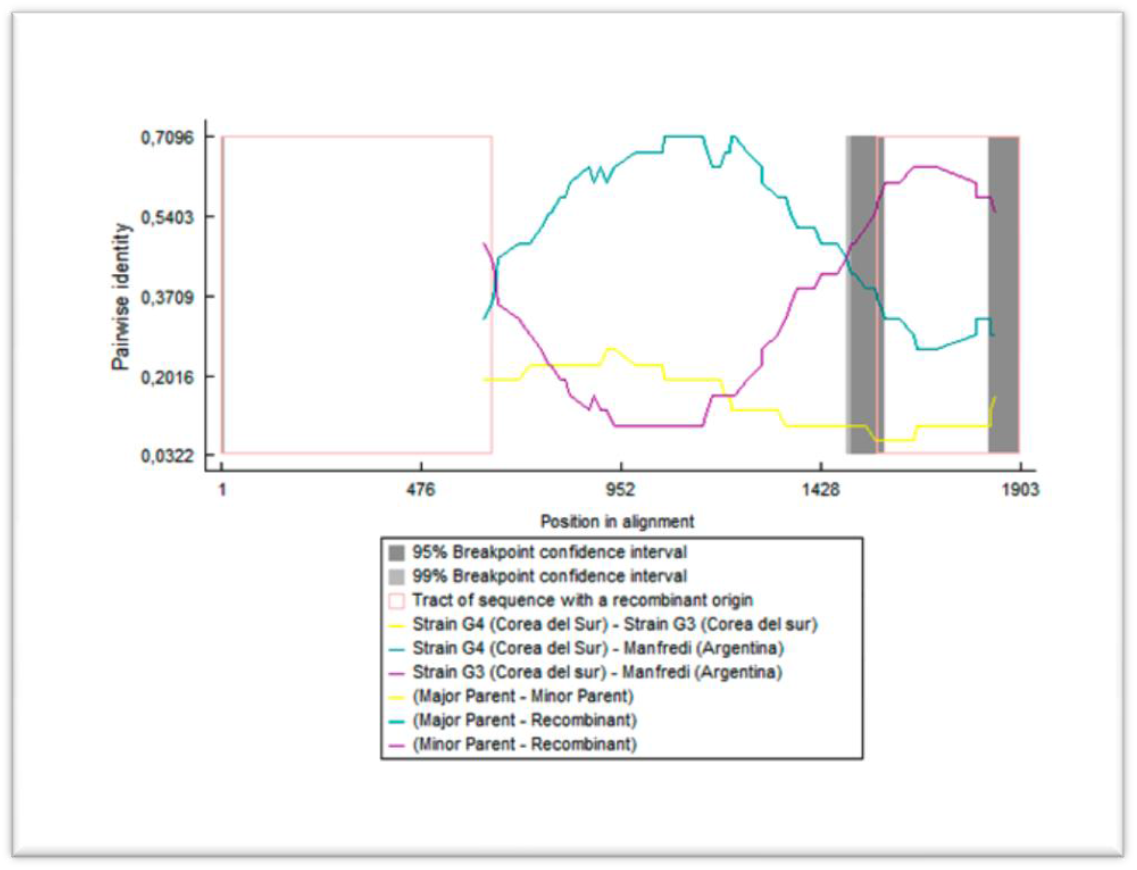
Recombination events detected in the CI segment of M isolate with parental LJZ010 (light blue) and G4 (violet); the corresponding breakpoints are included.

### 3. Early physiological alterations caused by the different isolates of SMV

Before the appearance of viral symptom (BS), SMV-NOA produced a differential behavior in ΦPII and MDA content in leaves with respect to the other isolates (Fig. 4b and F). Likewise, SMV-MJ caused a significant increase in total soluble sugar content with respect to the other isolates (Fig. 4d). All the SMV isolates induced a similar behavior in terms of Fv/FM, starch content and leaf area (Fig. 4a, c and e),

**Fig. 4.**
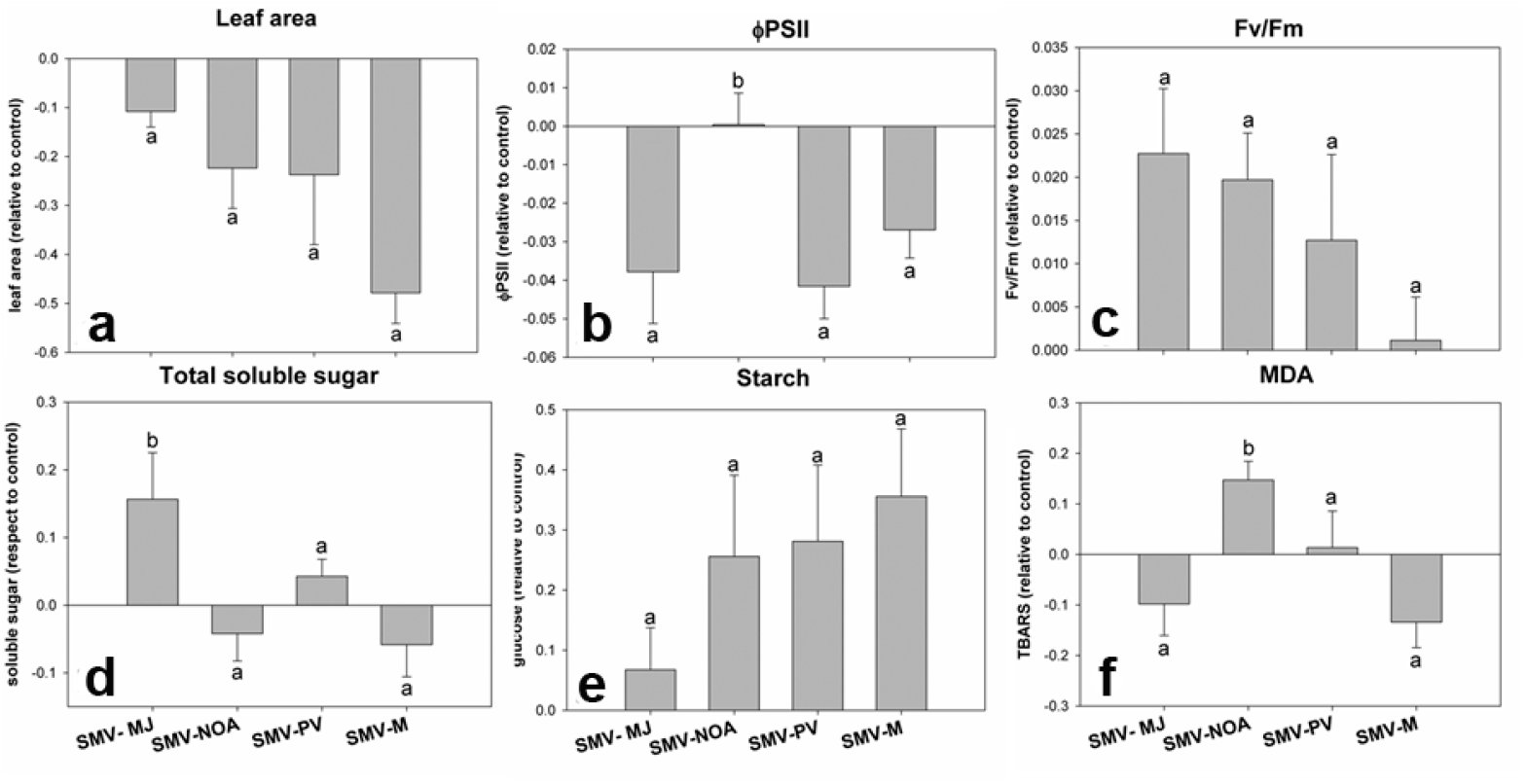
Early physiological alterations induced by different SMV isolates in soybean plants before macroscopic symptom appearance. **a.** leaf area; **b**. ΦPSII; **c.** Fv / Fm; **d.** Total soluble sugars; **e.** Starch; **f.** MDA. Sampling was carried out on the first trifoliate leaf 4 days after inoculation. Results are expressed as means ± SE of three independent experiments with at least three biological replicates each. Different letters indicate significant differences between treatments (DGC test. p <0.05).

### 4 Physiological alterations after viral symptom appearance

In order to analyze the effect of infection by the different virus isolates over time, we measured Fv/FM, starch content and leaf area in the same leaf, eight days after the early measurements.

SMV-NOA isolate caused an increase in MDA content, whereas (Fig. 5 f) SMV-M caused an increase in soluble sugar content, starch and MDA, without changes in ΦPSII (Fig. 5 b, d, e and f). SMV-PV isolate did not affect biomass production (Fig. 6). Moreover, relative viral concentration was measured after the appearance of symptoms, and no differences were observed among isolates (Supplementary Fig. 1).

**Fig. 5.**
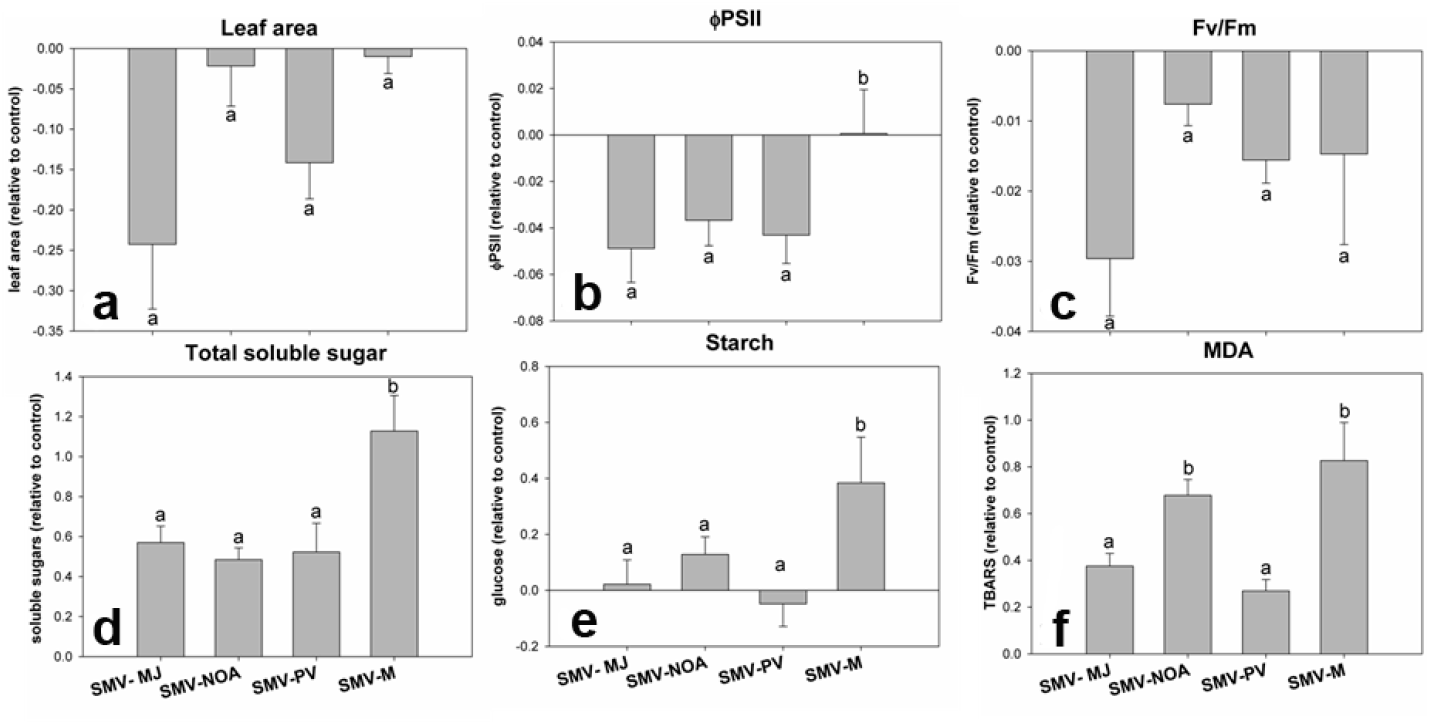
Physiological alterations induced by different isolates of SMV in soybean plants after the appearance of mosaic symptoms. **a**. leaf area; **b.** ΦPSII; **c**. Fv/Fm; **d**. Total soluble sugars; **e**. Starch; f. MDA. Sampling was carried out 12 days post-inoculation on the first trifoliate leaf. Results are expressed as means ± SE of three independent experiments with at least three biological replicates each. Different letters indicate significant differences between treatments (DGC test. p <0.05)

**Fig. 6.**
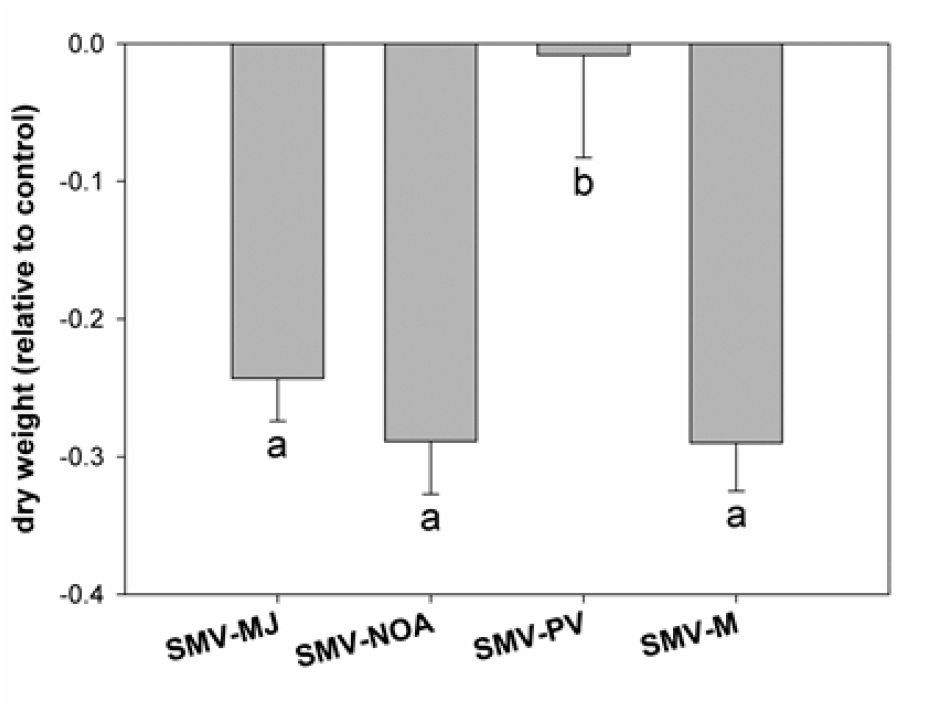
Dry weight of aboveground with respect to healthy control measured at 12 dpi (LS). Results are expressed as mean ± SE of three independent experiments with at least three biological replicates each. Different letters indicate significant differences between treatments (DGC test. p <0.05)

The SMV-MJ isolate was selected to study the response of two soybean cultivars susceptible to SMV infection, DM 4800 (DM) and Davis (D). Since the cultivar D showed lower infectivity than cultivar DM, physiological measurements were taken after the appearance of macroscopic symptoms (LS stage). Soybean cv Davies showed differential behavior with respect to DM in leaf area, ΦPSII, Fv/Fm and MDA (Fig. 7).

**Fig. 7.**
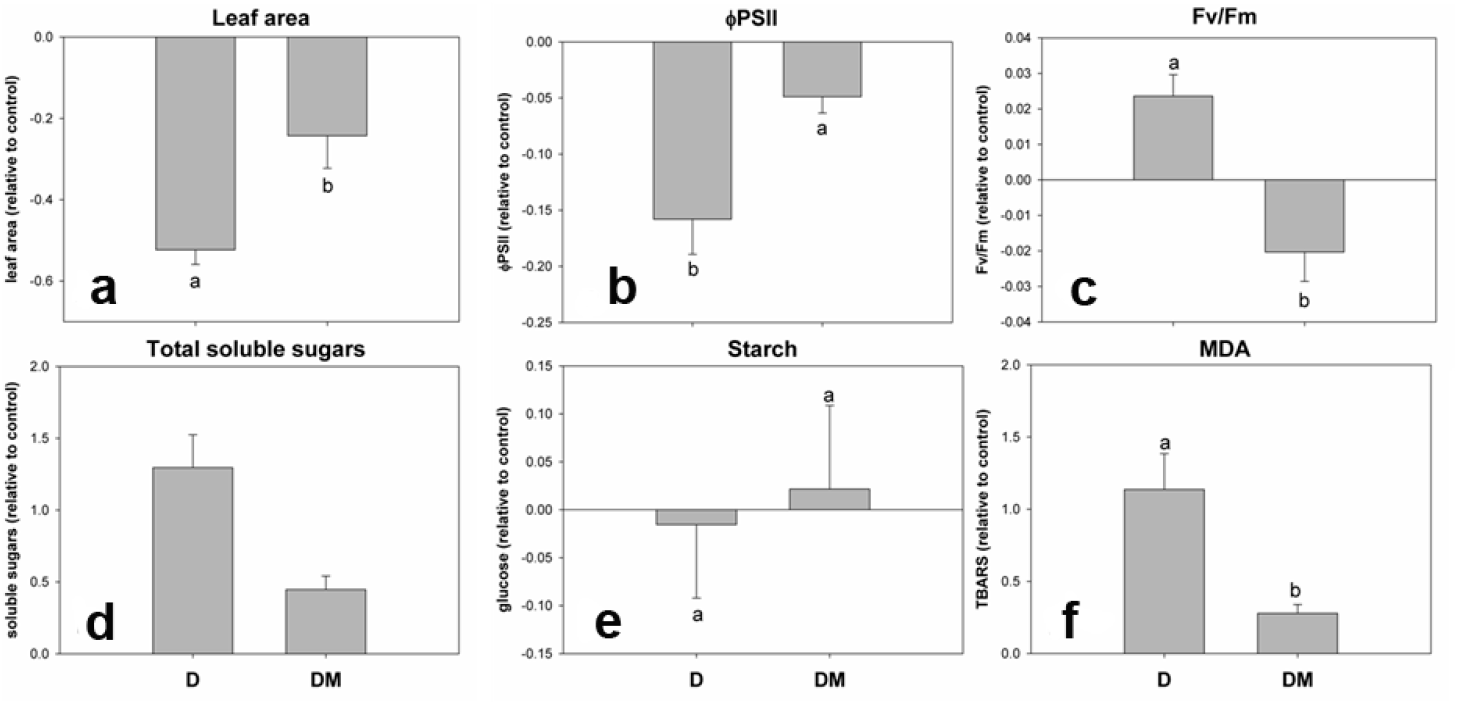
Physiological alterations induced by SMV in soybean plants of two susceptible cultivars after the appearance of mosaic symptoms. **a**. Leaf area; **b**. ΦPSII; **c**. Fv / FM; **d**. Total soluble sugars; **e.** Starch; **f.** MDA. Sampling was conducted 12 days after inoculation on the first trifoliate leaf. Results are expressed as ± SE of three independent experiments with at least three biological replicates each. Different letters indicate significant differences between treatments (DGC test. p <0.05)

## DISCUSSION

In this study, we performed the biological, molecular and physiological characterization of four SMV isolates, three geographical isolates (M, MJ and NOA) and one that causes symptoms of severe necrosis in some cultivars (PV). Although in the analyzed fragments SMV-PV isolate had high homology with SMV-NOA at the nucleotide level, it differed from the latter in pathogenicity, percentage of seed mottling, percentage of transmission by seeds and aphids, and plant physiological response. Specifically, those isolates showed significant differences in the effects on leaf area, ΦPSII and MDA content. The presence of the same recombination event in both isolates and their grouping in the phylogenetic trees suggest that they could belong to the same SMV race. However, we cannot rule out that the biological and physiological differences observed in both isolates could be explained by genetic differences in other parts of the genome. For instance, the interaction between the VPg protein (protein linked to the viral genome) and the plant transcription initiation factor, eIF4E, has been demonstrated to have an effect on the inhibition of gene expression during potyvirus infection due to destabilization of mRNA (Havelda et al., 2008).

So far, four independent loci for resistance to SMV have been identified (R*sv1*, R*sv3, Rsv4* and R*sv5*). In addition, multiple resistance alleles were reported for the loci R*sv1* and R*sv3* (Widyasari et al., 2020; Zheng et al., 2005). The emergence of resistance-breaking isolates can be attributed to the use of resistant cultivars, with a limited base of resistance to SMV subjected to selection pressure, due to mutations and/or recombination of the different virus strains (Choi et al., 2005). This may have been the case of SMV-PV isolate, since it produced a hypersensitive reaction in several cultivars, causing symptoms of necrosis in stem petioles and veins. In the pathogenicity tests, SMV-PV produced necrotic symptoms in all the evaluated cultivars, except for Buffalo and PI 483084, which contained R*sv1*-K and R*sv1*-h resistant genes. Proteins P3 and HC-Pro have been shown to be the effectors of resistance mediated by R*sv1*. In addition, the amino acids 823, 953 and 1112 of P3 were found to be important for the induction of the lethal systemic hypersensitive response (LSRV) (Hajimorad et al., 2006; R. H. Wen et al., 2013). Thus, SMV-PV could have emerged as a consequence of mutations in P3 and/or HC-Pro cistrons, which broke the resistance conferred, at least, by the alleles R*sv5, Rsv1 Rsv-1t* and *Rsv1k*. Thus, efforts should be made to complete the sequencing of these two cistrons with the aim to understand the biological differences between isolates, mainly considering that SMV-PV causes severe symptoms of necrosis.

CP and HC-Pro have been shown to play an important role in both aphid and seed transmission, whereas the P1 cistron is also a determinant of seed transmission (Jossey et al., 2013). The amino acid DAG sequence of the CP is conserved in most potyviruses and is involved in both types of transmission (seed and aphid). In addition, it has been shown that SMV induces seed coat mottling, presumably through the action of HC-Pro, which partially suppresses silencing of the Chalcone synthase (CHS) mRNAs (Atreya et al., 1990; Domier et al., 2003). With regards to the studied isolates, we found significant differences in transmission capacity, both by aphids and seeds. SMV-PV isolate showed the lowest aphid transmission capacity, and both SMV-PV and MJ isolates presented the lowest level of seed mottling, associated with a lower seed transmission capacity. However, the DAG motif found in the CP was present in all four isolates.

The analysis of the amino acid sequence of P1 showed that homology was greatest between the SMV-M and -MJ isolates and between SMV-PV and NOA isolates. The SMV-M isolate is the one that causes the highest percentage and severity of seed spotting, as well as the highest percentage of transmission by seeds and physiological differences at LS stage. It has been shown that SMV P1 protein interacts strongly with the Rieske Fe /S protein of soybean cytochrome b6f (Shi et al., 2007), an essential component of the electron transport chain in chloroplasts for photosynthesis. The interaction between chloroplast and the invading virus plays a critical role in viral infection and pathogenesis (Zhao *et al*., 2016). In this regard, this work, as previous works, showed a decrease in ΦPSII and a significant decrease in the CO_2_ fixation rate especially for SMV-MJ and PV (Andreola et al., 2019).Chloroplasts are the main source of intracellular reactive oxygen species (ROS) generation in green tissues, mainly under stress conditions. The virus ability to impair chloroplast function and disrupt the photosynthetic electron transport chain ultimately leads not only to the decrease of the carboxylation activity but also to ROS increase (Rodríguez *et al*. 2010; 2012; Zanini et al. 2021). Our results showed an increase in MDA content in soybean plants inoculated with SMV-NOA and SMV-M isolates at LS stage. MDA is a marker of oxidative lipid damage caused by stress (Arias et al., 2005). Moreover, SMV-NOA was the only isolate that produced an early increase in MDA content, without alteration of ΦPSII. These results suggest that oxidative damage measured through MDA in SMV-NOA infected plants before the onset of symptoms might be an early cellular oxidative process rather than the result of damage at the chloroplast level. Likewise, since SMV-NOA and -PV had high homology sequences in the analyzed fragment, we suggest that the physiological response at the chloroplast level could be related to other sequences that should be further explored.

Chloroplast alteration during SMV infection might be directly related to soluble sugar production. Present and previous results of our group have shown an increase in soluble sugars with SMV-MJ infection (Andreola et al., 2019). It is possible that the increase in soluble sugars observed in the compatible interaction between soybean and all SMV isolates is associated with the recycling of cellular components resulting from chloroplast damage. The accumulation of soluble sugars and the decrease in the ΦPSII and Fv/Fm suggest that the increase could be related to a greater import or lower export of sugars, or from intracellular recycling (Rodriguez et al., 2010; Andreola et al., 2019). These possible sugar sources are not mutually exclusive but might be operating in combination, and have to be explored in future studies.

SMV-MJ, one of the most severe isolates in terms of the reactions it produces in differential cultivars, showed early sugar alteration. On the other hand, the differential behavior of that virus isolate was found to occur not only between cultivars with different resistance (Cho and Goodman, 1979) or between susceptible and resistant cultivars (Arias et al., 2005), but also between susceptible cultivars. The soybean cv Davis (D) showed more severe symptoms than DM 4800 (DM) not only when infected with the SMV-MJ isolate in this work, but also when infected with the SMV-M isolate, which showed severe necrosis in the infected plants (unpublished data). Likewise, mechanical transmission had low efficiency in the SMV-M, SMV-NOA and SMV-PV isolates, suggesting a differential interaction of the same viral genome with different susceptible host plant genomes. In conclusion, knowing the physiological bases of viral infections and the mechanisms underlying plant infection by different races of viruses will contribute to the development of plants with tolerance to viral diseases.

## Acknowledgements

This work was supported by grants from the Agencia de Promoción Científica y Tecnológica, Argentina (PICT-2014-551), and Instituto Nacional de Tecnología Agropecuaria (INTA-). MR and NB are researchers of CONICET (Consejo Nacional de Investigaciones Científicas y Técnicas, Argentina). MR, NB and PRP are researchers of INTA.

## Compliance with ethical standards

### Conflict of interest

No potential conflicts of interest are disclosed.

### Research involving human participants and/or animals

The research involved neither human participants nor animals.

### Informed consent

not applicable.

---

**Supplementary Figure 1.**
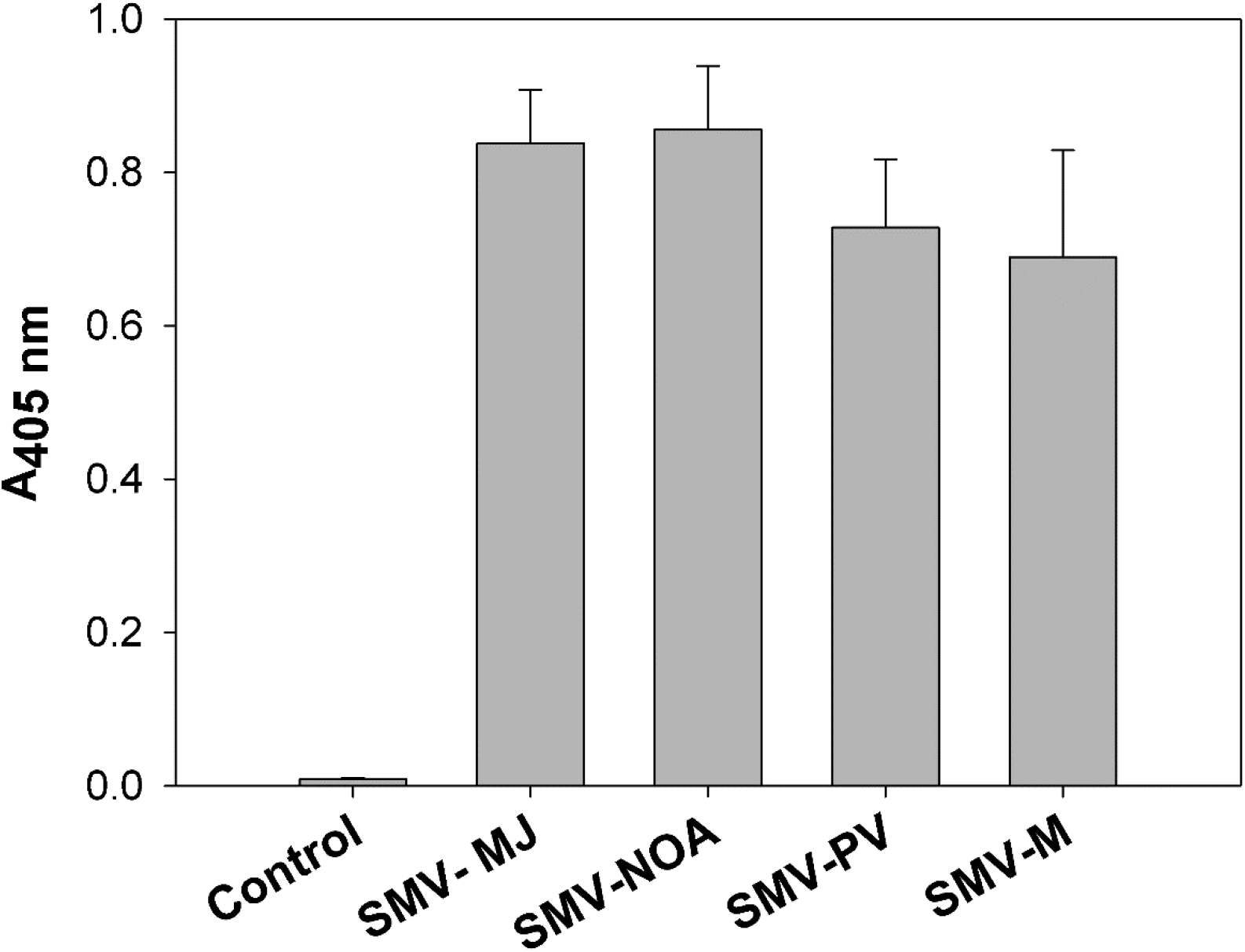
SMV accumulation (expressed as A 405 values of ELISA reactions) measured in the first trifoliate leaf at 12 days after inoculation (25μg of total protein. well^-1^). Results are expressed as mean ± SE of three independent experiments with at least three biological replicates each.

## Notes

### Competing Interest Statement

The authors have declared no competing interest.

